# AlphaPulldown – a Python package for protein-protein interaction screens using AlphaFold-Multimer

**DOI:** 10.1101/2022.08.05.502961

**Authors:** Dingquan Yu, Grzegorz Chojnowski, Maria Rosenthal, Jan Kosinski

## Abstract

**Summary:** The Artificial Intelligence-based structure prediction program AlphaFold-Multimer enabled structural modelling of protein complexes with unprecedented accuracy. Increasingly, AlphaFold-Multimer is also used to discover new protein-protein interactions. Here, we present *AlphaPulldown*, a Python package that streamlines protein-protein interaction screens and high-throughput modelling of higher-order oligomers using AlphaFold-Multimer. It provides a convenient command line interface, a variety of confidence scores, and a graphical analysis tool.

**Availability and implementation:** *AlphaPulldown* is freely available at https://www.embl-hamburg.de/AlphaPulldown.

## 1 Introduction

AlphaFold2 (Jumper *et al*., 2021) and AlphaFold-Multimer (Evans *et al*., 2022) have enabled structural modelling of monomeric proteins and protein complexes with accuracy comparable to experimental structures. Various modifications of AlphaFold2 have been developed to facilitate specific applications, such as ColabFold (Mirdita *et al*., 2022), which accelerates the program and exposes useful parameters. AlphaFold-Multimer can also be applied to screen large datasets of proteins for new protein-protein interactions (PPIs) (Humphreys *et al*., 2021; Bryant, Pozzati, and Elofsson, 2022) and to model combinations of proteins and their fragments when modelling complexes (Mosalaganti *et al*., 2022). To streamline such applications, we developed *AlphaPulldown* (Figure 1), a Python package that 1) provides a convenient command-line interface to run four typical scenarios in PPI screens and modelling of large complexes, 2) reduces the computing time by separating CPU- and GPU-based calculations, 3) allows selecting protein regions for modelling while retaining the original residue indexes, 4) provides a unique analysis pipeline that assesses the predicted interfaces with multiple scores and generates a Jupyter notebook for interactive analysis.

**Figure 1.**
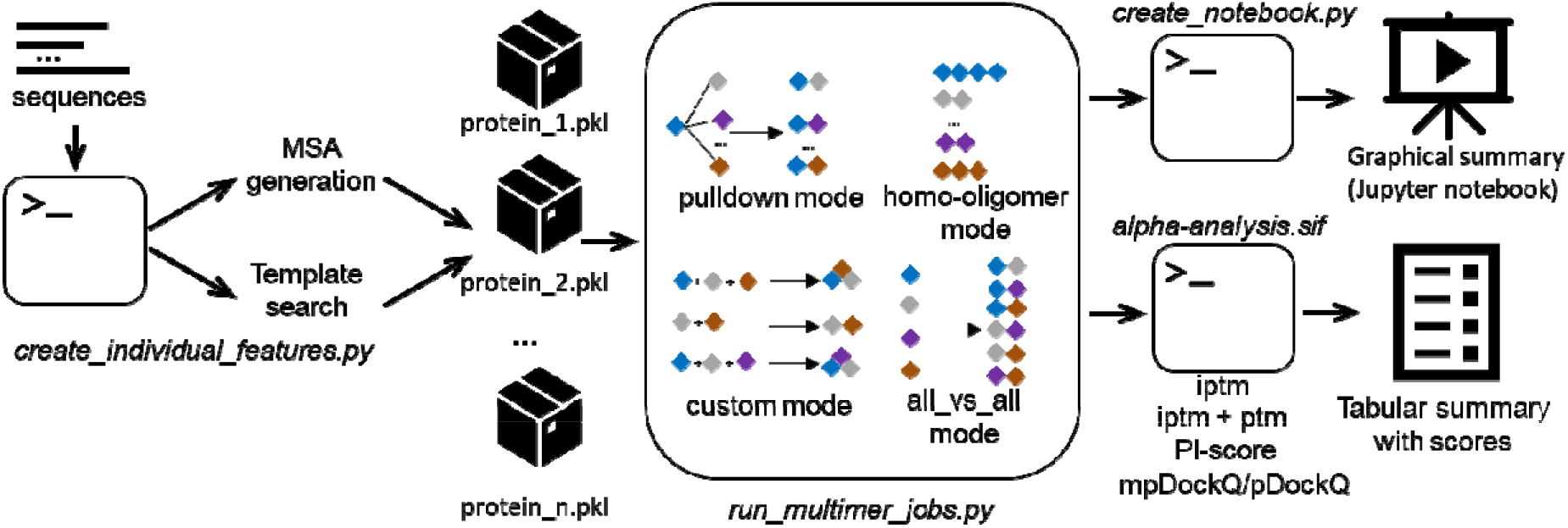
The workflow of *AlphaPulldown*. The first stage of *AlphaPulldown* runs on CPUs and is handled by the *create_individual_features.py* script, which takes protein sequences as input, generates MSAs, and finds structural templates for each protein. MSAs and structural features are stored in Python “pickle” files. The next stage takes these “pickle” files as input and predicts protein complex structures on GPUs. The user can choose from the following modes: 1) pulldown, 2) homooligomer, 3) custom, and 4) all versus all. Once all predictions are finished, *AlphaPulldown* generates a Jupyter notebook that provides a graphical summary and uses an analysis pipeline built in a Singularity image, *alpha-analysis.sif*, to produce a table with various model quality scores and interface properties.

## 2 Separation of stages

The original AlphaFold pipeline is composed of two main stages. The first uses CPUs to calculate multiple sequence alignments (MSAs) and to search for templates from PDB as input features for the second stage, which performs the actual structure prediction using GPUs. To speed up the pipeline, similar to other packages, *AlphaPulldown* separates these two stages, enabling the calculation of features for many sequences in parallel using multiple CPUs and prediction of structures using GPU.

## 3 Calculation of features

At this stage, *AlphaPulldown* searches each protein sequence against the original AlphaFold databases to pre-calculate MSA and identify template structures. It will also create features informing sequence by-organism pairing, which in the original AlphaFold-Multimer is performed only at the prediction state. *AlphaPulldown* stores these features in Python pickle files that can be reused for multiple PPI predictions.

## 4 Prediction modes

Once MSAs and structural template features are calculated, the user can choose from the following four modes of the main prediction.

### 4.1 Pull-down mode

Inspired by pulldown assays, this mode takes one or more proteins as “baits” and a list of other proteins as “candidates”. *AlphaPulldown* will automatically run AlphaFold-Multimer prediction between each bait and candidate. To demonstrate this mode, we show two examples. First, we screened for interactions of human eIF4G2 with proteins of the human translation pathway (Supplementary Note 1). The results confirmed a known interaction, revealed new interactions, and, based on the additional interface quality scores provided by *AlphaPulldown*, identified potential false positive and negative AlphaFold predictions (Supplementary Table 1 and Figure 1). Second, we modelled the interaction between the L and Z proteins of the Lassa virus, which could not be predicted using full-length sequences but only when screening Z against a series of fragments of L (Supplementary Note 2, Table 2, and Figure 2). This example shows that fragmenting large proteins may help AlphaFold find correct interaction interfaces. *AlphaPulldown* provides a convenient interface to specify any combination of residue ranges without needing to recalculate MSAs or template features. Moreover, *AlphaPulldown* keeps the residue numbering of original full-length sequences in the models.

### 4.2 All-versus-all mode

Based on a single list of proteins, *AlphaPulldown* will automatically generate all pairwise combinations and predict their structures. This mode can be useful to predict interaction networks and provide input for modelling large complexes with tools such as MoLPC (Bryant, Pozzati, Zhu, *et al*., 2022).

### 4.3 Homo-oligomer mode

This mode simplifies the process of modelling homo-oligomers and testing alternative homooligomeric states. The user only needs to provide an input file with desired oligomeric states, and *AlphaPulldown* will automatically run modelling for each state.

### 4.4 Custom mode

This mode allows the user to provide any combination of any number of proteins and their fragments as an input, not limited to pairwise predictions. This mode can be used, for example, to screen a pre-defined list of interactions from other sources such as crosslinking studies or model entire complexes. As mentioned above, even if fragments are used, the MSA and template features do not need to be recalculated and the resulting models keep the residue numbering of the fulllength sequences.

## 5 Analysis pipeline

For all models with an inter-chain predicted alignment error (PAE) below the user-defined cut-off, a Jupyter notebook will be generated to provide the user with a clear visual overview of these models as well as their ipTM and pLDDT scores (Supplementary Figure 3). The analysis pipeline will also return a CSV table containing the ipTM and ipTM+pTM scores reported by AlphaFold, a pDockQ score for dimers (Bryant, Pozzati, and Elofsson, 2022), mpDockQ score for multimers (Bryant, Pozzati, Zhu, *et al*., 2022), Protein Interface-score (PI-score)(Malhotra *et al*., 2021), and physical and geometrical properties calculated by the PI-score pipeline, such as the interface surface area or the number of hydrogen bonds (Supplementary Tables 1 and 2).

## 6 Conclusion

*AlphaPulldown* streamlines AlphaFold-multimer for PPI screens and modelling of large complexes fragment-by-fragment in a manner similar to that we applied to the human nuclear pore complex (Mosalaganti *et al*., 2022). It both facilitates the prediction process and provides a toolbox for a more thorough analysis of model confidence. We anticipate it will help the structural biology community unlock the full potential of AI-based structure prediction.

## Acknowledgements

We thank Agnieszka Obarska-Kosinska for testing the software and useful suggestions.

## Data availability

All the code and example data are available at https://www.embl-hamburg.de/AlphaPulldown and https://github.com/KosinskiLab/AlphaPulldown.

## References

Bryant, P., Pozzati, G., and Elofsson, A. (2022) Improved prediction of protein-protein interactions using AlphaFold2. Nature Communications 2022 13:1, 13, 1–11.

Bryant, P., Pozzati, G., Zhu, W., et al. (2022) Predicting the structure of large protein complexes using AlphaFold and sequential assembly. bioRxiv, 2022.03.12.484089.

Evans, R. et al. (2022) Protein complex prediction with AlphaFold-Multimer.

Humphreys, I. et al. (2021) Computed structures of core eukaryotic protein complexes. Science (1979), 374.

Jumper, J.et al. (2021) Highly accurate protein structure prediction with AlphaFold. Nature 2021 596:7873, 596, 583–589.

Malhotra, S. et al. (2021) Assessment of protein–protein interfaces in cryo-EM derived assemblies. Nature Communications 2021 12:1, 12, 1–12.

Mirdita, M. et al. (2022) ColabFold: making protein folding accessible to all. Nature Methods 2022 19:6, 19, 679–682.

Mosalaganti, S. et al. (2022) AI-based structure prediction empowers integrative structural analysis of human nuclear pores. Science (1979), 376.

