## Supplementary Materials for "AlphaPulldown – a Python package for protein-protein interaction screens using AlphaFold-Multimer"

**Supplementary Information**

**Supplementary Note 1**

eIF4G2 was chosen as a “bait” and the 294 proteins from human translation pathway retrieved from Reactome (StableID: R-HSA-72766.4) as “candidates”. The pulldown mode automatically created all 294 pairs between the “bait” and each “candidate”, and predicted their structures. Our screening yielded a high-quality model of eIF4G2 and eIF4A1 interaction (Uniprot: P60842, Supplementary Figure 1A), which has been previously identified by NMR and pulldown studies (IntAct: EBI-296519 and EBI-73449). The interaction obtained high ipTM+pTM, ipTM, pDockQ, PI-score, and favorable physical parameters (Supplementary Table 1). Similar scores were assigned to interaction with eIF4A2 (Uniprot: Q14240, Supplementary Figure 1B). Since both eIF4A1 and eIF4A2 form eIF4F complex variants with eIF4G1 (Complex Portal ComplexAc: CPX-5863 and CPX-5634, respectively), which is a paralog of eIF4G2, our screening result confirm that eIF4G2 might be able to substitute eIF4G1 and participate in eIF4F complex instead. An interaction with eIF3M (Uniprot: Q7L2H7) also obtained high scores, except a negative PI-score, likely due to a small interface (Supplementary Figure 1C). Notably, eIF4G2 can interact with eIF4A1 and eIF3M simultaneously in the context of 48S (PDB ID: 6ZMW). Thus, eIF4G2 might be able to replace eIF4G1 also in the 48S complex. An interaction with RPL41 (Uniprot: P62945) obtained very high scores from AlphaFold but low pDockQ and PI-score. The evaluation of the interface according to the physical parameters (Supplementary Table 1) reveals that interface is small and mediated through charged side chains (Supplementary Figure 1D, Supplementary Figure 1E). RPL41 is entirely engaged in binding RNA in the ribosome structures and thus likely this prediction is a false positive resulting from charge complementarity. Last, an interaction with SRP72 (Uniprot: O76094) received poor AlphaFold scores but high pDockQ and PI-score, and large interface area (Supplementary Figure 1F). Thus, while speculative, this prediction might represent a false negative prediction by AlphaFold and a good candidate for experimental testing. Overall, the AlphaPulldown results for eIF4G2 show a typical scenario the users might encounter and demonstrate how false positive and false negative predictions could be detected with our analysis tool.

| jobs | Num_intf_residues | Polar | Hydrophobhic | Charged | contact_pairs | sc | hb | sb | int_solv_en | int_area | pi_score | iptm+ptm | mpDockQ/pDockQ | iptm |
|---|---|---|---|---|---|---|---|---|---|---|---|---|---|---|
| P78344_and_P60842 | 54 | 0.259 | 0.259 | 0.333 | 49 | 0.576 | 18 | 19 | -10.40 | 2535.35 | 1.13 | 0.7700 | 0.7420 | 0.8278 |
| P78344_and_Q7L2H7 | 15 | 0.067 | 0.200 | 0.333 | 10 | 0.489 | 6 | 5 | -1.90 | 735.79 | -0.75 | 0.7680 | 0.7416 | 0.8190 |
| P78344_and_P62945 | 2 | 0.000 | 0.500 | 0.500 | 1 | 0.389 | 4 | 9 | 2.67 | 685.21 | -0.28 | 0.7463 | 0.3968 | 0.8170 |
| P78344_and_Q14240 | 55 | 0.255 | 0.273 | 0.327 | 47 | 0.565 | 22 | 27 | -5.68 | 3002.93 | 1.50 | 0.7501 | 0.7420 | 0.8041 |
| P78344_and_P62913 | 9 | 0.111 | 0.222 | 0.444 | 5 | 0.477 | 4 | 2 | -0.62 | 758.07 | -0.78 | 0.6896 | 0.7268 | 0.7561 |
| P78344_and_P41091 | 6 | 0.333 | 0.000 | 0.500 | 6 | 0.520 | 3 | 0 | 0.31 | 484.04 | 0.30 | 0.6923 | 0.6371 | 0.7343 |
| P78344_and_P46778 | 12 | 0.000 | 0.500 | 0.333 | 11 | 0.449 | 7 | 16 | -3.63 | 929.08 | 0.02 | 0.6551 | 0.7390 | 0.7013 |
| P78344_and_P62857 | 25 | 0.120 | 0.400 | 0.200 | 24 | 0.341 | 5 | 4 | -3.25 | 1138.99 | -1.87 | 0.6219 | 0.7306 | 0.6729 |
| P78344_and_Q07020 | 5 | 0.200 | 0.000 | 0.600 | 3 | 0.276 | 3 | 10 | 4.23 | 683.35 | -0.44 | 0.6379 | 0.6996 | 0.6724 |
| P78344_and_P60866 | 16 | 0.250 | 0.313 | 0.375 | 18 | 0.445 | 9 | 8 | -4.40 | 1007.69 | -0.88 | 0.6192 | 0.7388 | 0.6652 |
| P78344_and_P43897 | 21 | 0.143 | 0.381 | 0.095 | 21 | 0.450 | 4 | 3 | -5.09 | 812.32 | -0.73 | 0.6184 | 0.7158 | 0.6473 |
| P78344_and_P49207 | 20 | 0.150 | 0.400 | 0.250 | 17 | 0.371 | 5 | 8 | -4.84 | 1411.45 | -1.93 | 0.5952 | 0.7390 | 0.6199 |
| P78344_and_P15880 | 14 | 0.286 | 0.357 | 0.286 | 15 | 0.491 | 3 | 1 | -7.94 | 424.87 | 0.33 | 0.5669 | 0.7180 | 0.6123 |
| P78344_and_Q92901 | 7 | 0.286 | 0.143 | 0.286 | 7 | 0.507 | 5 | 1 | -1.58 | 404.03 | -0.48 | 0.5871 | 0.4825 | 0.6093 |
| P78344_and_Q9Y291 | 14 | 0.143 | 0.143 | 0.500 | 15 | 0.272 | 2 | 7 | -0.27 | 786.97 | -1.70 | 0.5663 | 0.6724 | 0.6031 |
| P78344_and_Q7Z2W9 | 38 | 0.105 | 0.289 | 0.526 | 62 | 0.288 | 10 | 11 | -8.26 | 1286.54 | -1.24 | 0.5535 | 0.7399 | 0.5955 |
| P78344_and_P60228 | 24 | 0.125 | 0.375 | 0.250 | 29 | 0.404 | 7 | 3 | -10.98 | 1168.50 | -0.05 | 0.5422 | 0.7361 | 0.5801 |
| P78344_and_O76094 | 53 | 0.358 | 0.302 | 0.151 | 54 | 0.536 | 25 | 2 | -26.75 | 2188.93 | 1.08 | 0.5422 | 0.7413 | 0.5788 |
| P78344_and_P62495 | 12 | 0.167 | 0.250 | 0.500 | 11 | 0.319 | 8 | 14 | 0.76 | 1368.37 | -1.31 | 0.5347 | 0.7381 | 0.5497 |
| P78344_and_P26641 | 14 | 0.214 | 0.357 | 0.286 | 11 | 0.415 | 4 | 3 | -2.58 | 911.94 | -2.13 | 0.3827 | 0.7378 | 0.3833 |
| P78344_and_P82912 | 10 | 0.100 | 0.200 | 0.600 | 13 | 0.406 | 6 | 12 | -0.81 | 739.57 | -0.94 | 0.3854 | 0.6714 | 0.3775 |
| P78344_and_Q9BYN8 | 8 | 0.000 | 0.750 | 0.250 | 8 | 0.596 | 1 | 5 | -9.28 | 561.91 | 0.38 | 0.3600 | 0.6900 | 0.3548 |
| P78344_and_P23588 | 29 | 0.414 | 0.345 | 0.207 | 23 | 0.565 | 8 | 5 | -17.45 | 1547.37 | 1.54 | 0.3228 | 0.7353 | 0.3203 |
| P78344_and_P62851 | 11 | 0.000 | 0.273 | 0.727 | 14 | 0.524 | 7 | 2 | -6.03 | 785.19 | 0.50 | 0.2975 | 0.6870 | 0.2654 |

Supplement Table 1 Screening results of eIF4G1 (Uniprot: P78344) against human translation initiation pathway. Only predictions with at least one PAE value between chains lower than 5 are shown. The table is ordered by iPTM score. iPTM and iPTM+pTM scores are reported by AlphaFold. mpDockQ/pDockQ is calculated using the formula given by (Bryant, Pozzati, and Elofsson, 2022; Bryant, Pozzati, Zhu, *et al.*, 2022). pi_score and the rest of the columns are reported by PI-score pipeline(Malhotra *et al.*, 2021). Num_intf_residues: number of residues at the interface. Polar: number of polar residues (Ser, Thr, Asn, Gln, His and Tyr) at the interface. Hydrophobic: number of hydrophobic residues (Ala, Leu, Ile, Val, Phe, Trp, Cys, Met) at the interface. Charged: number of charged residues (Asp, Glu, Lys, Arg) at the interface. contact_pairs: number of atomic contacts between the interface residues. sc: geometric shape complementarity of protein-protein interfaces. sc ranges between 0 and 1 with sc=1 being two proteins mesh precisely. hb: number of hydrogen-bonds in the interface. sb: number of salt-bridges at the interface. int_solv_en: interface solvation energy. int_area: interface surface area that will be inaccessible to solvent upon the interface formation.

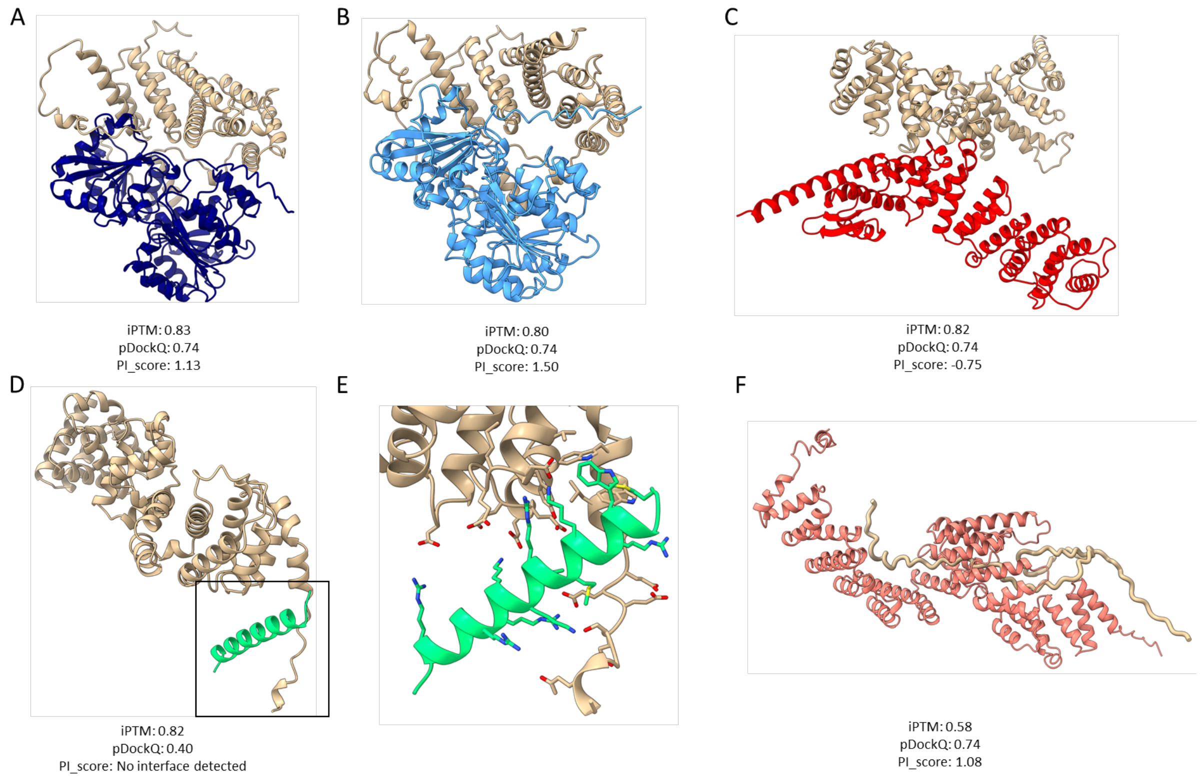


**Supplementary Figure 1:** Example results of the predicted eIF4G2 interaction screen. EIF4G2 is colored brown, and the interacting partner in another distinct color. The non-interacting domains of eIF4G2 and flexible disordered regions are hidden for clarity. **(A)** Interaction with eIF4A1 (Uniprot: P60842). **(B)** Interaction with eIF4A2 (Uniprot: Q14240). **(C)** Interaction with eIF3M (Uniprot: Q7L2H7). **(D)** Potentially false positive interaction with RPL41 (Uniprot: P62945). **(E)** Enlarged view of the predicted interface with RPL41. **(F)** Potentially false negative interaction with SRP72 (Uniprot: O76094).

### Supplementary Note 2

Lassa L protein is known to interact with its Z protein(Loureiro *et al.*, 2011). L has more than 2200 residues, and when modelling full-length L with Z, neither *AlphaPulldown* nor default AlphaFold Multimer found any interface, likely owing to the large size of L. Therefore, we split L into five different fragments, according to its structural domains(Kouba *et al.*, 2021), and screened for possible interactions against Z protein. Eventually, *AlphaPulldown* successfully found an interface between Z and a fragment of L between residue 290 and 1398, which is in agreement with an experimental structure of the L-Z complex (Xu *et al.*, 2021).

| jobs | Num_intf_residues | Polar | Hydrophobhic | Charged | contact_pairs | sc | hb | sb | int_solv_en | int_area | pi_score | iptm+ptm | mpDockQ/pDockQ | iptm |
|---|---|---|---|---|---|---|---|---|---|---|---|---|---|---|
| O73557_and_O09705 | None | None | None | None | None | None | None | None | None | None | No interface detected | 0.5492 | 0.0609 | 0.4872 |
| O73557_and_O09705_290-1398 | 12 | 0.250 | 0.500 | 0.083 | 13 | 0.446 | 4 | 0 | -13.19 | 651.00 | -0.18 | 0.6060 | 0.7355 | 0.5406 |
| O73557_and_O09705_1830-2217 | 11 | 0.545 | 0.182 | 0 | 10 | 0.169 | 0 | 0 | -10.45 | 685.53 | -1.46 | 0.2656 | 0.7077 | 0.1839 |
| O73557_and_O09705_196-297_700-1398 | None | None | None | None | None | None | None | None | None | None | No interface detected | 0.2755 | 0.0183 | 0.1463 |
| O73557_and_O09705_1398-1830 | None | None | None | None | None | None | None | None | None | None | No interface detected | 0.2464 | 0.0367 | 0.1239 |
| O73557_and_O09705_1-200 | None | None | None | None | None | None | None | None | None | None | No interface detected | 0.2203 | 0.2107 | 0.1202 |

Supplement Table 2: Screening results of Lassa virus Z protein (Uniprot: O73557) against full-length and fragments of L protein (Uniprot: O09705). Results with full-length L protein are listed at the top. Results with L protein fragments are ordered by iPTM score. iPTM and iPTM+pTM scores are reported by AlphaFold. mpDockQ/pDockQ is calculated using the formula given by(Bryant, Pozzati, and Elofsson, 2022; Bryant, Pozzati, Zhu, *et al.*, 2022). pi_score and the rest of the columns are reported by PI-score pipeline(Malhotra *et al.*, 2021). Num_intf_residues: number of residues at the interface. Polar: number of polar residues (Ser, Thr, Asn, Gln, His and Tyr) at the interface. Hydrophobic: number of hydrophobic residues (Ala, Leu, Ile, Val, Phe, Trp, Cys, Met) at the interface. Charged: number of charged residues (Asp, Glu, Lys, Arg) at the interface. contact_pairs: number of atomic contacts between the interface residues. sc: geometric shape complementarity of protein-protein interfaces. sc ranges between 0 and 1 with sc=1 being two proteins mesh precisely. hb: number of hydrogen-bonds in the interface. sb: number of salt-bridges at the interface. int_solv_en: interface solvation energy. int_area: interface surface area that will be inaccessible to solvent upon the interface formation.

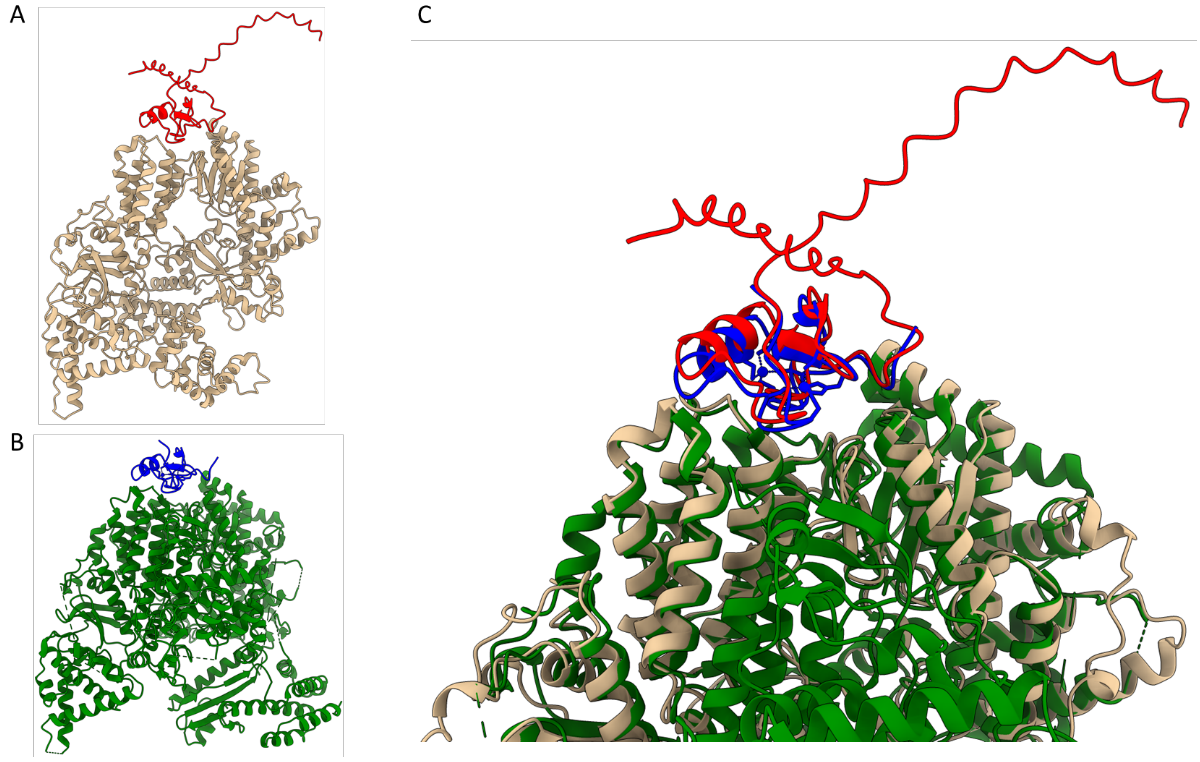


**Supplementary Figure 2**: Comparison between Lassa virus L protein fragment interacting with Z protein predicted by *AlphaPulldown* vs cryo-EM structure of L-Z protein complex (PDB: 7ELA). **(A)** *AlphaPulldown* modelled a region of L protein between residue 290 and 1398 (brown) and the Z protein is predicted to be interacting with this region in the position shown in red. **(B)** Cryo-EM structure of L protein (green) and Z protein (blue). **(C)** Superposition of our model and the experimental structure shows that the interface between L and Z protein in our model agrees with the experimental data.

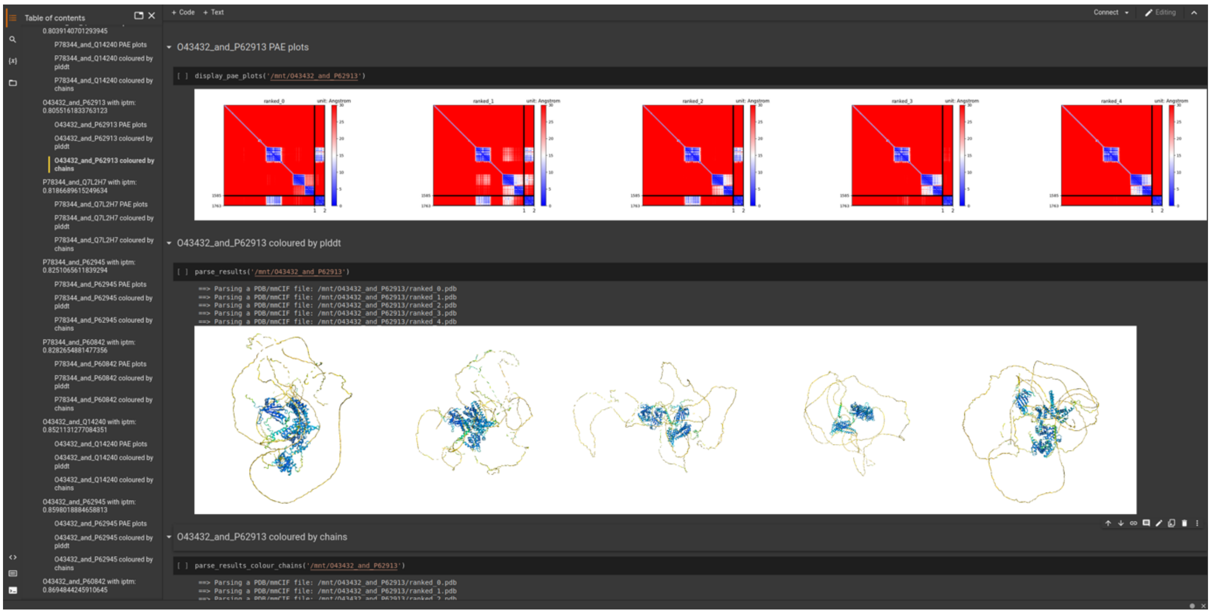


**Supplementary Figure 3**: A screenshot of an example Jupyter notebook generated by *AlphaPulldown.* On the left, individual jobs are listed as bookmarks. When the user clicks one job's bookmark and executes corresponding cells, the notebook will automatically display 1) PAE plots 2) predicted model colored by pLDDT scores of each residue, and 3) predicted structure colored by chains.

### Supplementary references

Kouba,T. *et al.* (2021) Conformational changes in Lassa virus L protein associated with promoter binding and RNA synthesis activity. *Nat. Commun.*, **12**, 1–18.

Loureiro,M.E. *et al.* (2011) Molecular Determinants of Arenavirus Z Protein Homo-Oligomerization and L Polymerase Binding. *J. Virol.*, **85**, 12304–12314.

Xu,X. *et al.* (2021) Cryo-EM structures of Lassa and Machupo virus polymerases complexed with cognate regulatory Z proteins identify targets for antivirals. *Nat. Microbiol. 2021 67*, **6**, 921–931.
